# Genomic SEM Provides Insights into the Multivariate Genetic Architecture of Complex Traits

**DOI:** 10.1101/305029

**Authors:** Andrew D. Grotzinger, Mijke Rhemtulla, Ronald de Vlaming, Stuart J. Ritchie, Travis T. Mallard, W. David Hill, Hill F. Ip, Andrew M. McIntosh, Ian J. Deary, Philipp D. Koellinger, K. Paige Harden, Michel G. Nivard, Elliot M. Tucker-Drob

**Affiliations:** Department of Psychology, University of Texas at Austin, Austin, TX USA; Department of Psychology, University of California, Davis, Davis, CA USA; Department of Economics, Vrije Universiteit Amsterdam, Amsterdam, the Netherlands; Erasmus University Rotterdam Institute for Behavior and Biology, Rotterdam, the Netherlands; Centre for Cognitive Ageing and Cognitive Epidemiology, University of Edinburgh, Edinburgh, UK; Department of Psychology, University of Edinburgh, Edinburgh, UK; Department of Biological Psychology, VU University Amsterdam, Amsterdam, the Netherlands; Division of Psychiatry, University of Edinburgh, Edinburgh, UK; Population Research Center, University of Texas at Austin, Austin, TX USA

**Author notes:** These authors jointly directed this work (Michel G. Nivard, Elliot M. Tucker-Drob). Correspondence to Andrew D. Grotzinger.

## Abstract

Methods for using GWAS to estimate genetic correlations between pairwise combinations of traits have produced “atlases” of genetic architecture. Genetic atlases reveal pervasive pleiotropy, and genome-wide significant loci are often shared across different phenotypes. We introduce genomic structural equation modeling (Genomic SEM), a multivariate method for analyzing the joint genetic architectures of complex traits. Using formal methods for modeling covariance structure, Genomic SEM synthesizes genetic correlations and SNP-heritabilities inferred from GWAS summary statistics of individual traits from samples with varying and unknown degrees of overlap. Genomic SEM can be used to identify variants with effects on general dimensions of cross-trait liability, boost power for discovery, and calculate more predictive polygenic scores. Finally, Genomic SEM can be used to identify loci that cause divergence between traits, aiding the search for what uniquely differentiates highly correlated phenotypes. We demonstrate several applications of Genomic SEM, including a joint analysis of GWAS summary statistics from five genetically correlated psychiatric traits. We identify 27 independent SNPs not previously identified in the univariate GWASs, 5 of which have been reported in other published GWASs of the included traits. Polygenic scores derived from Genomic SEM consistently outperform polygenic scores derived from GWASs of the individual traits. Genomic SEM is flexible, open ended, and allows for continuous innovations in how multivariate genetic architecture is modeled.

## Genomic Structural Equation Modeling

Genome-wide association studies (GWASs) are rapidly identifying loci affecting multiple phenotypes.^1,2^ Moreover, using cross-trait versions of methods such as genomic-relatedness-based restricted maximum-likelihood (GREML)^3^ and LD-score regression (LDSC)^4^ researchers have identified genetic correlations between diverse traits, *e.g*., polycystic ovary syndrome and pubertal timing,^5^ amyotrophic lateral sclerosis (ALS) and schizophrenia,^6^ and anorexia nervosa and obsessive-compulsive disorder.^7^ More generally, these analyses are suggestive of constellations of phenotypes affected by shared sources of genetic liability, but they do not permit the causes of the observed genetic correlations to be investigated systematically. Here we introduce Genomic Structural Equation Modeling (Genomic SEM), a new method for modeling the multivariate genetic architecture of constellations of traits and incorporating genetic covariance structure into multivariate GWAS discovery. Genomic SEM formally models the genetic covariance structure of GWAS summary statistics from samples of varying and potentially unknown degrees of overlap, in contrast to methods^8^ that model phenotypic covariance structure using raw data. Moreover, Genomic SEM allows the user to specify and compare a range of different hypothesized multivariate genetic architectures, which improves upon existing approaches for combining information across genetically correlated traits to aid in discovery.^9^

One powerful feature of Genomic SEM is the capability to model shared genetic architecture across phenotypes with factors that may be treated as broad genetic liabilities. This allows researchers to identify variants with effects on general dimensions of cross-trait liability, boost power for discovery, and calculate more valid and predictive polygenic scores. Genomic SEM can also evaluate whether the multivariate genetic architecture implied by a specific model is applicable at the level of individual variants using developed estimates of heterogeneity. When certain SNPs only influence a subset of genetically correlated traits, a key assumption of other multivariate approaches is violated.^9^ SNPs with high heterogeneity estimates can be flagged as likely to confer disproportionate or specific liability toward individual traits or disorders, can be removed when constructing polygenic risk scores, or studied specifically to understand the nature of heterogeneity. These heterogeneity estimates act as safeguards against false inference when considering a locus specific to one trait in its effect on a set of correlated traits.

We validate key properties of Genomic SEM with a series of simulations and illustrate the flexibility and utility of Genomic SEM with several analyses of real data. These include a joint analysis of GWAS summary statistics from five genetically correlated psychiatric case-control traits: schizophrenia, bipolar disorder, major depressive disorder (MDD), post-traumatic stress disorder (PTSD), and anxiety. We model genetic covariances among the traits using a general factor of psychopathology (*p*), for which we identify 27 independent SNPs not previously identified in the univariate GWASs, 5 of which can be validated based on separate GWASs. Polygenic scores derived using this *p*-factor consistently outperform polygenic scores derived from GWASs of the individual traits in out-of-sample prediction of psychiatric symptoms. Other demonstrations include a multivariate GWAS of neuroticism items, an exploratory factor analysis of anthropometric traits, and a simultaneous analysis of the unique genetic associations between schizophrenia, bipolar disorder, and educational attainment.

## Results

Genomic SEM is a Two-Stage Structural Equation Modeling approach.^10-12^ In Stage 1, the empirical genetic covariance matrix and its associated sampling covariance matrix are estimated. The diagonal elements of the sampling covariance matrix are squared standard errors (*SEs*). The off-diagonal elements index the extent to which sampling errors of the estimates are associated, as may be the case when there is sample overlap across GWAS. In principle, these matrices may be obtained using a variety of methods for estimating SNP heritability, their genetic covariance, and their *SEs*. Here we use a novel version of LDSC that accounts for potentially unknown degrees of sample overlap by populating the off-diagonal elements of the sampling covariance matrix. The same strengths, as well as assumptions and limitations that are known to apply to LDSC^13,14^ apply to its extension used here and to Genomic SEM. In Stage 2, a SEM is estimated by minimizing the discrepancy between the model-implied genetic covariance matrix and the empirical covariance matrix obtained in the previous stage. We highlight results from weighted least squares (WLS) estimation that weights the discrepancy function using the inverse of the diagonal elements of the sampling covariance matrix, and produces model *SEs* using the full sampling covariance matrix. In the **Online Supplement**, we report results from an alternative normal theory maximum likelihood (ML) estimation method. We evaluate fit with the standardized root mean square residual (SRMR), model χ^2^, Akaike Information Criteria (AIC), and Comparative Fit Index (CFI; **Online Method**).^11,15^

Genomic SEM can be employed as a tool for multivariate GWAS based on univariate summary statistics. First, the genetic covariance matrix and its associated sampling covariance matrix are expanded to include SNP effects. A Genomic SEM is then specified in which SNP effects occur at the level of a latent genetic factor defined by several phenotypes, at the level of the genetic component of each of several (potentially genetically correlated) phenotypes, or some combination of the two. The Genomic SEM is then run once per SNP (or each set of SNPs, should the user incorporate multiple SNPs into a model) to obtain its effects within the multivariate system.

We provide an index that quantifies the extent to which an observed vector—consisting of univariate regression effects of a given SNP on each of the phenotypes—can be explained by a *common pathway* model that assumes that the effects on the phenotypes are entirely mediated by the common genetic factor(s). In other words, the index enables the identification of loci that do and do not operate on the individual phenotypes exclusively by way of their associations with the common factor(s). Because of its intuitive and mathematical similarity to the meta-analytic Q-statistic used in standard meta-analyses to index heterogeneity of effect sizes^16^ we label this heterogeneity statistic, Q_SNP_. Q_SNP_ is a χ^2^-distributed test statistic with larger values indexing a violation of the null hypothesis that the SNP acts entirely through the common factor(s) (**Online Method**).

## Validation via Simulation

We performed 100 runs of Genomic SEM on raw individual-level genotype data for which we simulated multivariate phenotypic data to conform to a single genetic factor model (a latent trait that partially causes 5 observed outcomes). Across the 100 simulations Genomic SEM estimates closely matched the parameters specified in the generating population (Supplementary Fig. 1). Model *SEs* also closely matched the standard deviations of parameter estimates. We also compared fit statistics (CFI, AIC, and model χ^2^) for the correctly specified common factor model and two deliberately misspecified models: (i) a model in which all indicators were constrained to have the same factor loading, and (ii) a model for which the loading of the third indicator was set to 0. As expected, results indicated that the common factor model matching the generating population was favored ≥ 99% of the time across model fit indices (Supplementary Fig. 2).

### Confirmatory Factor Analysis of Genetic Covariance Matrices

We provide two examples of confirmatory factor analysis (CFA) using Genomic SEM. We specifically factor analyze a genetic covariance and sampling covariance matrix with estimates populated from multivariable LDSC for psychiatric traits and for indicators of neuroticism. Recent findings indicate that the comorbidity across psychiatric disorders is captured by a latent, general psychopathology factor that is known as the *p*-factor and is widely supported based on previous results.^17,18^ We tested for the presence of a genetic *p*-factor using Genomic SEM with European-only summary statistics for schizophrenia, bipolar disorder, major depressive disorder (MDD), post-traumatic stress disorder (PTSD), and anxiety (Table S1 for phenotypes and sample sizes). Model fit was adequate (χ^2^(5) = 89.55, AIC = 109.5, CFI = .848, SRMR = .212).^1^ [1 For large samples, model χ^2^, such as those observed in GWAS, is expected to be highly significant and is more appropriate in Genomic SEM as a means of comparing nested models.] Results indicated that schizophrenia and bipolar disorder loaded the strongest onto the genetic *p*-factor (Supplementary Fig. 3), a pattern of findings that closely replicates prior phenotypic findings.^19^ We next tested for the presence of a single neuroticism factor using summary statistics from 12 item-level indicators from UK Biobank (UKB; Table S1) as estimated using the Hail software.^20^ Model fit was good (χ^2^(54) = 4884.10, AIC =4932.10, CFI = .893, SRMR = .109). Results revealed a common neuroticism factor with strong positive loadings for all indicators (Supplementary Fig. 4).

### Exploratory Factor Analysis of a Genetic Covariance Matrix

Exploratory factor analysis (EFA) was applied to the genetic covariance matrix for nine anthropometric traits from the EGG and GIANT consortia (Table S1). A heatmap of the genetic correlation matrix suggests two primary factors that index overweight and early life-growth phenotypes (Supplementary Fig. 5). Results of the EFA indicated that two factors explained 60% of the total genetic variance. A follow-up CFA (Supplementary Fig. 6) within Genomic SEM was specified based on the EFA parameter estimates (standardized loadings > .2 were retained). The CFA showed good fit to the data (χ^2^(25) = 10397.45, AIC = 10437.45, CFI = .951, SRMR = .089). Results indicated highly significant loadings, and a small correlation between the two factors (*r_g_* = .11, *SE* = .03, *p* < .001). This indicates that early life physical growth is modestly associated with later life obesity traits via genetic pathways.

### Genetic Multivariable Regression (Replicating GWIS)

Nieuwboer et al. (2016)^21^ use summary statistics for educational achievement (EA)^22^ and both schizophrenia and bipolar disorder^23^ to determine if genetic correlations with EA are driven by variation specific to either disorder. EA is genetically correlated with schizophrenia (*r_g_* = .148, *SE* = .050, *p* = .003) and bipolar disorder (*r_g_* = .273, *SE* = .067, *p* < .001). Using a method called genome-wide inferred statistics (GWIS), they find that the correlation of EA with schizophrenia unique of bipolar is small (*r_g_* = .040, *SE* = .082, *p* = .627), whereas the genetic correlation between bipolar unique of schizophrenia and EA is far less attenuated (*r_g_* = .218, *SE* = .102, *p* = .032). We use Genomic SEM with the aim of replicating these results using a conceptually similar, but statistically distinct, framework. We present this example to demonstrate that Genomic SEM is not limited to factor analytic models, but can be used to construct and test an array of hypotheses using a general SEM approach.

Using the same univariate GWAS summary statistics employed in the original application of GWIS, we used Genomic SEM to fit a structural multivariable regression model in which the genetic component of EA was simultaneously regressed onto the genetic components of schizophrenia and bipolar disorder. Fit indices are not reported as this was a fully saturated model (i.e., *df* = 0). Results confirmed the findings by Nieuwboer et al. (2016);^21^ the conditional standardized association between schizophrenia and EA was quite small (*b_g_* = -.016, *SE* = .096, *p* = .867), whereas there was a strong conditional standardized association between bipolar disorder and EA (*b_g_* = .283, *SE* = .113, *p* = .012; Supplementary Fig. 7).

### SNP Effects

A powerful application of Genomic SEM is to include individual SNP effects in both the genetic covariance matrix and the sampling covariance matrix, in order to estimate the effect of a given SNP on the latent genetic factor(s). If the summary statistics are composed of *M* different SNPs, then *M* models are estimated to obtain genome-wide summary statistics for the latent factor. We apply this method to the *p*-factor and neuroticism models presented above. LD-independent hits are defined here as *r*^2^ < .1 in a 500-kb window, with the exception of a 1-mb window for chromosomes 6 and 8. The effect of 128 independent loci crossed the threshold for genome wide significance for the *p*-factor (*p* < 5 × 10^-8^; Supplementary Figs. 8-9 for QQ-plot and Manhattan plot of univariate estimates; Figs. 1 and 2 for factor model and Manhattan plots). Of the 128 loci, 27 independent loci were not previously identified in any of the univariate GWAS (Table 1, Table S2). Of these 27 loci, five were identified as either genome wide significant or suggestive of significance (*p* < 1 × 10^-5^) in an outside study of an overlapping trait. The effect of 118 loci crossed the threshold for genome wide significance for neuroticism, with 38 loci not identified in the univariate item-level GWASs (Table S3). Intercepts from LDSC analyses of the summary statistics produced by Genomic SEM indicated that results were not due to uncontrolled inflation for either the *p*-factor (intercept = .987, *SE* = .014) or neuroticism (intercept = .997, *SE* = .001).

**Fig. 1.**
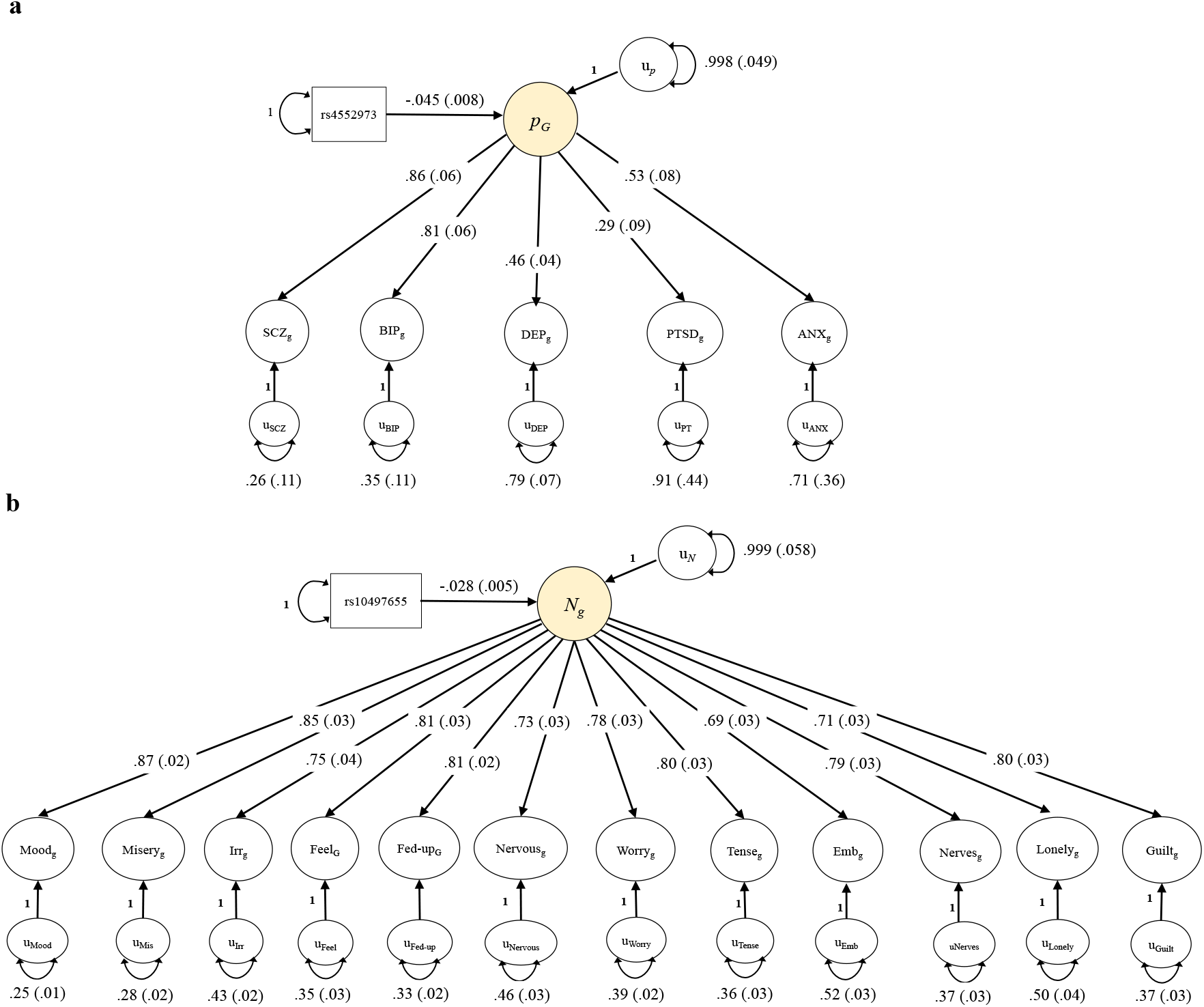
Genomic SEM solutions for *p*-factor and neuroticism factor models with SNP effect. Standardized results from using Genomic SEM (with WLS estimation) to construct a genetically defined *p*-factor of psychopathology (panel a) and a genetic neuroticism factor (panel b) with a lead independent SNP predicting the factors. *SEs* are shown in parentheses. For a model that was standardized with respect to the outcomes only, the effect of the SNP was -.093 (*SE* = .017; SNP variance = .252) for the *p*-factor, and for neuroticism the SNP effect was -.042 (*SE* = .007, SNP variance = .432); this can be interpreted as the expected standard deviation unit difference in the latent factor per effect allele. SCZ = schizophrenia; BIP = bipolar disorder; DEP = major depressive disorder; PTSD = post-traumatic stress disorder; ANX = anxiety. Irr = irritability; Feel = sensitivity/hurt feelings; fed-up = fed-up feelings; emb = worry too long after embarrassment.

**Fig 2.**
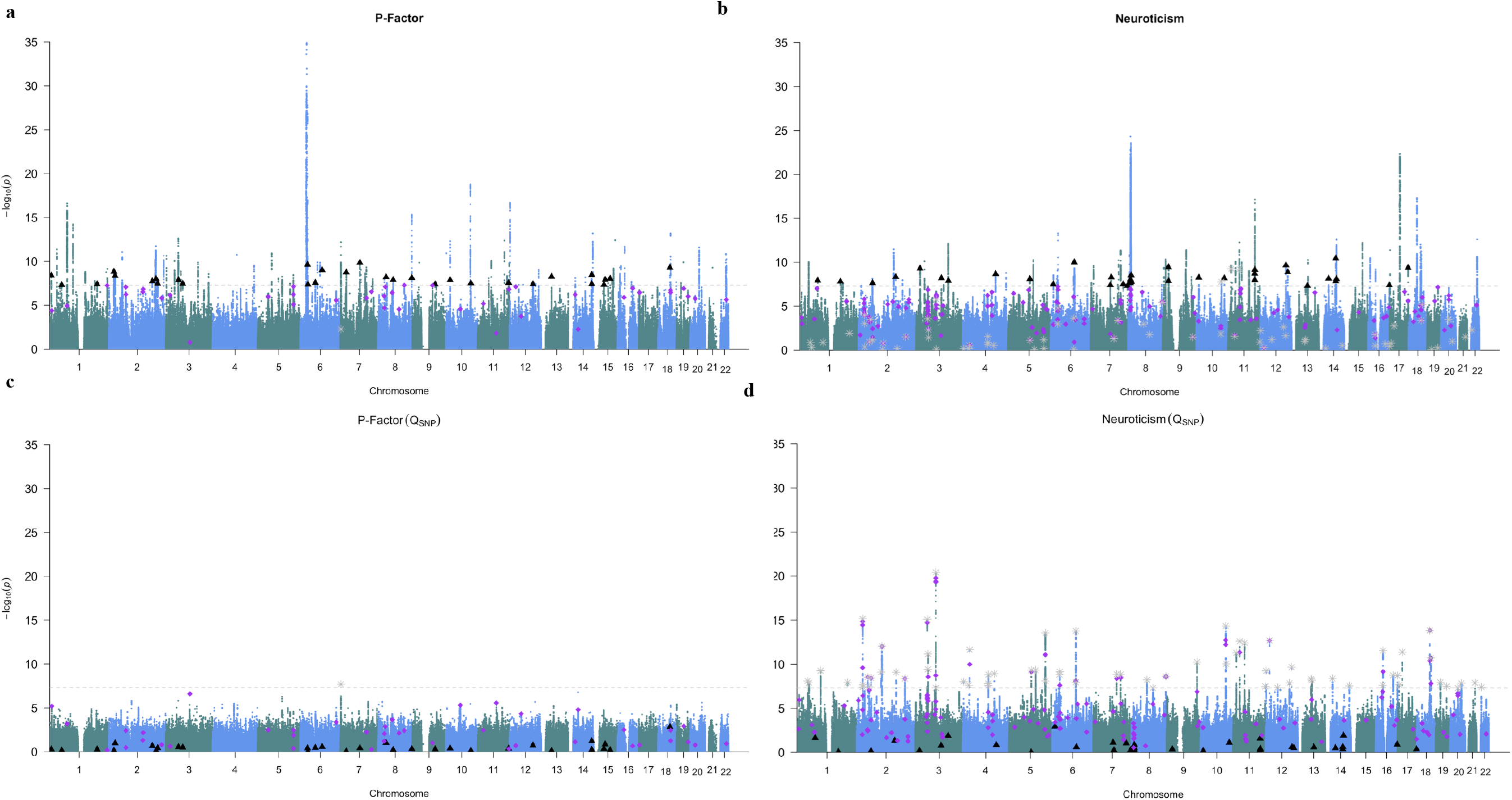
Manhattan plots of unique, independent hits from Genomic SEM. Genomic SEM (with WLS estimation) was used to conduct multivariate GWASs of the *p*-factor (panel a) and neuroticism (panel b). Manhattan plots are shown for SNP effects (left panels) and for Q_SNP_ (right panels). The gray dashed line marks the threshold for genome wide significance (*p* < 5 × 10^-8^). In all four panels, black triangles denote independent hits for SNP effects from the GWAS of the general factor that were not in LD with independent hits for the univariate GWAS or hits for Q_SNP_. In all four panels, purple diamonds denote independent hits for the SNP effects from univariate GWASs that were not in LD with independent hits from the GWAS of the general factor. Grey stars denote independent hits for Q_SNP_.

**Table 1.** Summary of multivariate (Genomic SEM) and univariate GWAS results.

| | Lead SNPs<br>($p < 5 \times 10^{-8}$) | $Q_{\text{SNP}}$ hits | Unique<br>Hits | No. of<br>gene sets | No.<br>prioritized<br>genes | No.<br>tissues<br>and cells | Mean<br>$\chi^2$ |
| --- | --- | --- | --- | --- | --- | --- | --- |
| **P-Factor** |  |  |  |  |  |  |  |
| Genomic<br>SEM (WLS) | 128 | 1 (1) | 31 (4) | 71 | 37 | 24 | 1.88 |
| Schizophrenia | 127 | - | 34 (0) | 2 | 25 | 21 | 1.82 |
| Bipolar | 4 | - | 4 (0) | 0 | 0 | 0 | 1.15 |
| MDD | 5 | - | 5 (0) | 0 | 0 | 0 | 1.31 |
| PTSD | 0 | - | 0 (0) | 0 | 0 | 0 | 1.01 |
| Anxiety | 1 | - | 1 (0) | 0 | 0 | 0 | 1.03 |
| **Neuroticism** |  |  |  |  |  |  |  |
| Genomic<br>SEM (WLS) | 118 | 69 (6) | 38 (0) | 1 | 19 | 20 | 1.64 |
| Mood | 43 | - | 19 (5) | 0 | 0 | 15 | 1.37 |
| Misery | 31 | - | 6 (4) | 0 | 0 | 0 | 1.32 |
| Irritability | 36 | - | 17 (4) | 0 | 0 | 0 | 1.37 |
| Hurt Feelings | 24 | - | 11 (0) | 0 | 0 | 0 | 1.33 |
| Fed-up | 38 | - | 21 (6) | 0 | 0 | 0 | 1.36 |
| Nervous | 41 | - | 25 (12) | 0 | 0 | 0 | 1.36 |
| Worry | 56 | - | 26 (6) | 0 | 13 | 0 | 1.46 |
| Tense | 19 | - | 10 (3) | 0 | 0 | 0 | 1.32 |
| Embarrass | 17 | - | 6 (2) | 0 | 0 | 0 | 1.33 |
| Nerves | 12 | - | 7 (3) | 0 | 0 | 0 | 1.26 |
| Lonely | 6 | - | 4 (3) | 0 | 0 | 0 | 1.19 |
| Guilt | 21 | - | 8 (1) | 0 | 0 | 0 | 1.28 |

*Note*. In parentheses for Q_SNP_ reports how many Q_SNP_ hits were in LD with hits identified as significant for the common factor. Unique hits for the common factor refers to lead SNPs that were not in LD with hits for the individual indicators. Unique hits for the individual indicators refers to hits for the respective indicator that were not in LD with hits for the common factor. Unique hits for the common factor excluded hits in LD with Q_SNP_ hits. For unique hits for indicators, values in parentheses indicate whether any of these hits were identified as significant for Q_SNP_. For unique hits for the common factor, values in parentheses indicate hits that were in LD with previously reported indicator hits that were removed due to missing values across the other indicators. The single Q_SNP_ hit for WLS estimation of the *p*-factor was significant for both the common factor and schizophrenia. For the common factor and the indicators, independent hits were defined using a pruning window of 500 kb and *r*^2^ > 0.1. For chromosomes 6 and 8, an additional pruning filter was used of 1,000,000 kb and *r*^2^ > 0.1 to account for long-range LD due to the MHC region and pericentric inversion, respectively. For univariate statistics, we used only the SNPs present across all indicators in order to facilitate a direct comparison to Genomic SEM results.

#### General Trends

Mean χ^2^ statistics were higher for the Genomic SEM-derived summary statistics of common factors relative to univariate indicators (Table 1). It is important to note here that, whereas Genomic SEM may boost power in many cases, this is not the primary purpose of the method. Rather, it is to identify the relation between SNPs and observed phenotypes as meditated through a user-specified model and to concurrently evaluate the construct validity of said model. Inspecting the distribution of univariate *p*-values for the newly identified SNPs for the general factors indicated that these SNPs were generally characterized by relatively low *p*-values, albeit not low enough to cross the genome-wide significance threshold for any individual phenotype (Supplementary Figs. 10-11).

#### Q_SNP_ Results

Results revealed 1 and 69 independent Q_SNP_ loci for the *p*-factor and neuroticism, respectively (Fig. 2; Supplementary Fig. 12 for QQ-plot). For neuroticism, significant Q_SNP_ estimates were obtained for SNPs that were highly significant for some traits but not others (Table S4; Supplementary Fig. 13). The association between *p*-values for SNP effects and Q_SNP_ estimates were minimal (Supplementary Fig. 14). Comparing the Q_SNP_ estimates for these SNPs with those that were identified as significant for only the *p*-factor or neuroticism indicated that SNPs not identified for the common factors were characterized, as would be expected, by larger Q_SNP_ estimates (i.e., greater heterogeneity in individual effects; Supplementary Fig. 15). Intercepts from LDSC analyses of the Q statistics also indicated that results for Q_SNP_ were not attributable to inflation (*p*-factor: intercept = .978, *SE* = .009; neuroticism: intercept = .963, *SE* = .009). Slopes from these LDSC analyses indicated genetic signal in heterogeneity (*p*-factor: *Z* = 13.65, *p*-value = 6.68E-42; neuroticism: *Z* = 30.23, *p*-value = 9.98E-201).

#### Performance under controlled missingness

We contrast estimates obtained from the common factor model of neuroticism described above with estimates for a GWAS with an imposed missing structure. We first transformed the binary scale neuroticism items into a smaller number of quantitative scores. To do so, we created three parcels of neuroticism items consisting of 4 items each with scores ranging from 0 to 4, at which point it is appropriate to treat the parcel as continuous.^24^ Parcels were constructed based on factor loadings obtained using EFA of the genetic covariance matrix of the neuroticism items (Table S5). Of the 300,000 participants, 100,000 non-overlapping participants were removed from two of the three parcels for missing data models. The most well-powered results were for Genomic SEM of the individual neuroticism items presented above, indicating that construction of composite indices via averaging, though convenient, removes multivariate information that can otherwise be retained with Genomic SEM (Table S6). Genomic SEM analyses that incorporated supplemental information from parcels containing imposed missing data consistently outperformed GWAS of individual parcels with complete data, and performed nearly as well as analyses of complete data across all three parcels.

#### Parcel Comparison of Q_SNP_

Using the three constructed parcels without any missing data, the distribution of *p*-values was compared across SNPs with high (*p* < 5e-8) and low (*p* > 5e-3) Q_SNP_ estimates from the item-level Genomic SEM analysis of neuroticism and that that were genome wide significant hits in at least one of the parcels. These results indicated that, for SNPs with a higher Q_SNP_ for the common factor, there was more discordance of effect sizes among three lower-order factors relative to SNPs that produced lower heterogeneity estimates (Supplementary Fig. 16). The average difference between the highest and lowest −log10 *p*-values was 10.56 and 4.96 for high and low Q_SNP_, respectively. This suggests that Q_SNP_ is appropriately indexing discordance in SNP level effects across genetically correlated indicators.

#### Polygenic Prediction

We re-estimated the *p*-factor model using the summary statistics from the SCZ and MDD GWASs that did not overlap with the UKB dataset, in order to predict psychiatric symptoms in UKB (Supplementary Fig. 17 for phenotypic model). We compared the magnitude of out-of-sample-prediction for the *p*-factor PGSs predicting the phenotypic *p*-factor and individual psychiatric domains relative to the prediction using PGSs derived from univariate summary statistics (Fig. 3, Table S7). The PGSs for the genetic *p*-factor predicted more variance in depression, psychotic experiences, mania, anxiety, PTSD and a phenotypic *p*-factor than any univariate PGS.

**Fig. 3.**
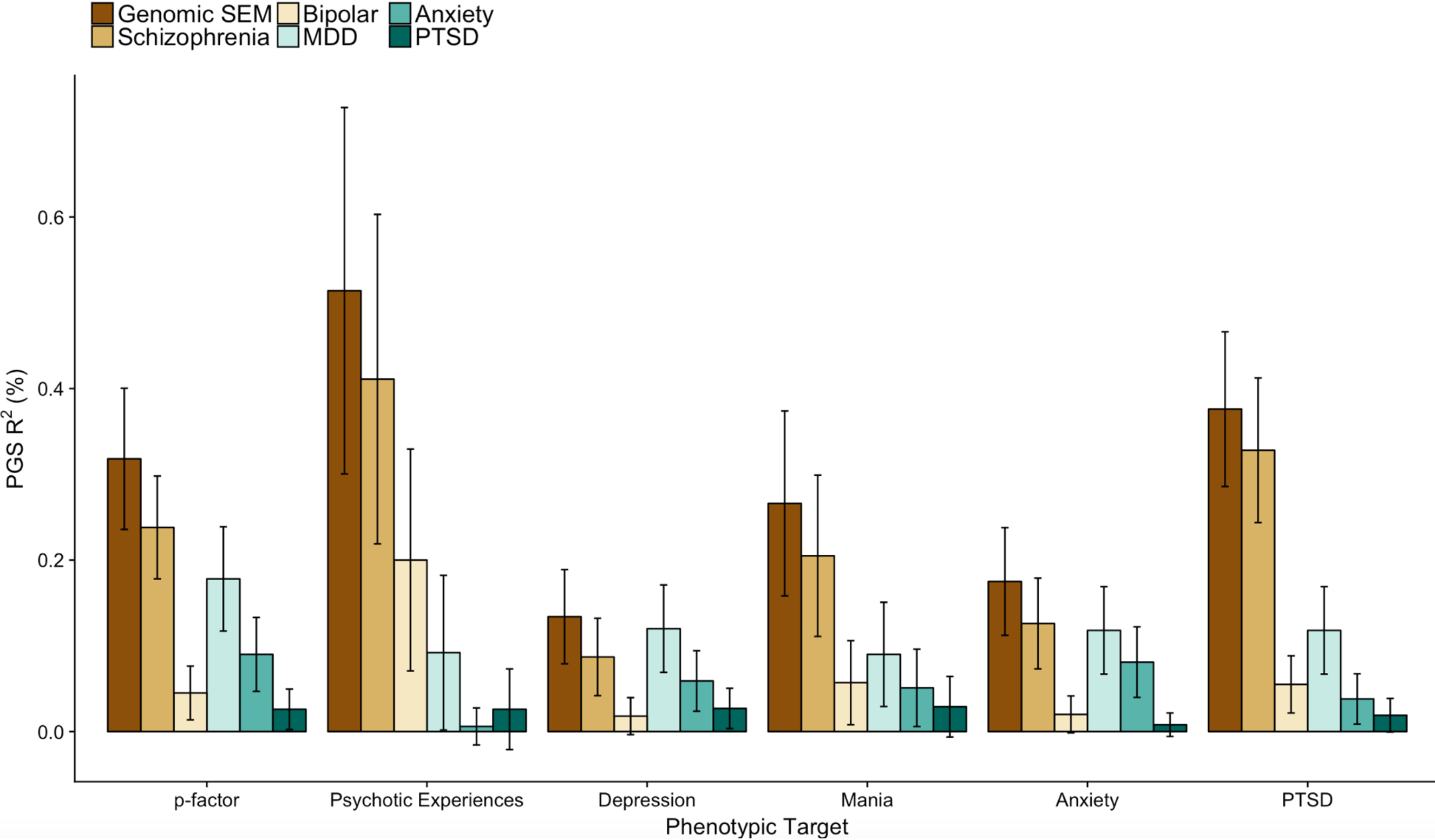
Out-of-sample prediction using Genomic SEM based and univariate based polygenic scores. Polygenic scores (PGSs) were constructed using the same set of SNPs for all predictors. R^2^ (%) on the y-axis indicates the percentage of variance (possible range: 0-100) explained in the outcome unique of covariates. The summary statistics for Genomic SEM were estimated using WLS. Error bars indicate 95% confidence intervals estimated using the delta method. Phenotypes were constructed for European participants in the UKB for five symptom domains and for a general *p* factor spanning all five symptom domains.

#### Biological Annotation

The biological function of the SNPs related to the *p*-factor and neuroticism was examined using DEPICT.^25^ Table 1 presents the number of enriched gene sets, prioritized genes, and enriched tissues and cell types across the univariate statistics and common factors (Supplementary Tables S8-S16 for detailed output). Common factors produced far more informative results than the individual indicators. As expected, all of the tissue enrichment for the common factors was identified in the nervous system (Fig. 4). Neuroticism prioritized genes indicated a central role of synaptic activity (e.g., *STX1B, NR4A2, PCLO*), including glutamatergic neurotransmission (*GRM3).* The *p*-factor gene sets were largely characterized by communication between neurons (e.g., “dendrite development”, “dendritic spine”, “abnormal excitatory postsynaptic potential”). Biological annotation of Q_SNP_ statistics for neuroticism indicated that genes within the 69 loci related to neuroticism, but not through a single factor, include: *GRIA1*, a glutamate receptor subunit (i.e. involved in signaling is excitatory neurons) which has previously been related to schizophrenia,^26^ chronotype,^27^ and autism;^28^ and *PCDH17*, a gene involved in cellular connections in the brain that has been related to intelligence.^29^

**Fig. 4.**
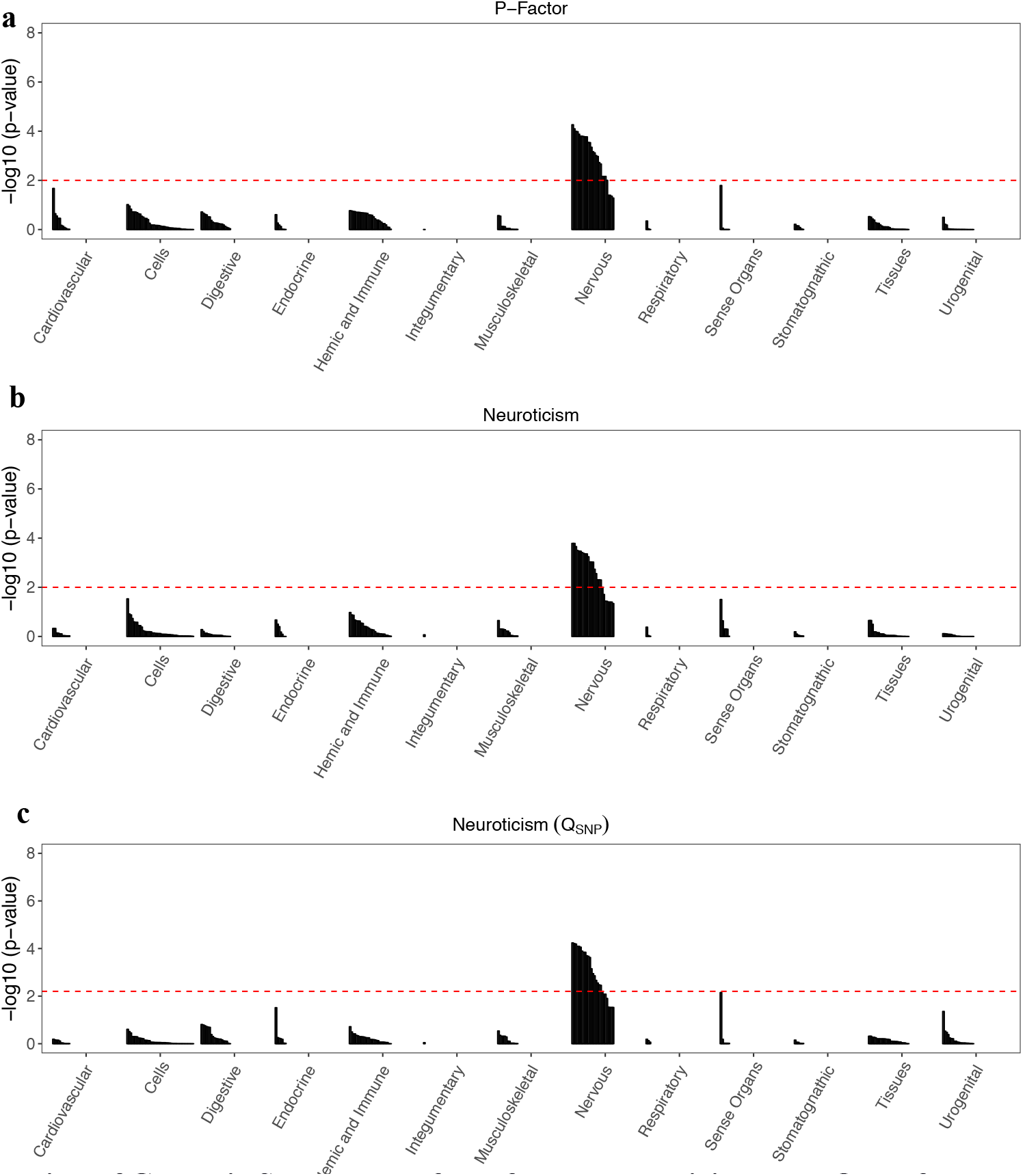
Biological annotation of Genomic SEM results for *p*-factor, neuroticism, and Q_SNP_ of neuroticism. Results from tissue enrichment analyses conducted using DEPICT based on Genomic SEM results for the *p*-factor (panel a), neuroticism (panel b) and Q_SNP_ estimation for neuroticism (panel c) using WLS estimation. The red, dashed line indicates the false discovery rate at .05. As expected, the majority of enriched tissues were in the nervous system for both common factors and Q_SNP_ estimate.

## Discussion

Genomic SEM is a novel method for modeling the multivariate genetic architecture of constellations of genetically correlated traits and incorporating genetic covariance structure into multivariate GWAS discovery. In contrast to methods^8^ that model phenotypic, rather than genetic covariance structure, and rely on raw data, Genomic SEM employs summary GWAS data to model genetic covariance structure. Genomic SEM is computationally efficient, accounts for potentially unknown degrees of sample overlap, and allows for flexible specification of covariance structure, such that several broad classes of structured covariance models can be applied. The Genomic SEM approach shares benefits of some existing approaches^9^ for boosting power by combining information across genetically correlated phenotypes. However, Genomic SEM uniquely allows one to compare different hypothesized genetic covariance architectures and to incorporate such architectures into multivariate discovery. Importantly, shared genetic liabilities across phenotypes can be explicitly modeled as factors that may be treated as broad genetic risk factors with equally broad downstream consequences.

In contrast to approaches that assume homogeneity of effects across SNPs,^9^ Genomic SEM includes diagnostic indices for its key assumptions, including a test for heterogeneity, Q_SNP_, that can be applied at the level of the individual SNPs. This offers the unique ability to identify SNPs that confer specific risk to individual disorders, symptoms, or indicators. This question may be of particular interest as the large degrees of genetic overlap identified across phenotypes (e.g., bipolar disorder and schizophrenia) beg the question: what are the genetic causes of phenotypic divergence? Whereas previous GWASs have combined items tapping genetically-related phenotypes into a single score, or even combined cases with different diagnoses to obtain a shared genetic effect, Genomic SEM allows researchers to interrogate shared genetic effects between diagnoses or indicators, while concurrently testing for causes of divergence (i.e., loci that are related only to a specific phenotype, or subset of phenotypes, but not the more general liability). In the context of neuroticism, for example, we identified 69 loci that were significantly involved in one manifestation of neuroticism but whose effects were not shared through a common factor, offering novel evidence of biological heterogeneity in the etiology of a construct long thought to be unidimensional. Because Genomic SEM relies only on GWAS summary data, it can be applied to a broad spectrum of traits, including gene expression, hormones, metabolites, brain structure and functions, behaviors, and psychiatric disorders and medical diseases.

## Acknowledgements

Elliot M. Tucker-Drob, K. Paige Harden, and Andrew D. Grotzinger, were supported by NIH Grant R01HD083613. Elliot M. Tucker-Drob, Stuart J. Ritchie, and Ian J. Deary were supported by NIH Grant R01AG054628. Elliot M. Tucker-Drob and K. Paige Harden were each supported by Jacobs Foundation research fellowships. Elliot Tucker-Drob and K. Paige Harden are members of the Population Research Center at the University of Texas at Austin, which is supported by NIH grant R24HD042849. Michel G. Nivard is supported by a Royal Netherlands Academy of Science Professor Award to Dorret I. Boomsma (PAH/6635), ZonMw grant:“ Genetics as a research tool: A natural experiment to elucidate the causal effects of social mobility on health.” (pnr: 531003014) and ZonMw project: “Can sex - and gender-specific gene expression and epigenetics explain sex-differences in disease prevalence and etiology?” (pnr:849200011). Hill F Ip is supported by the “Aggression in Children: Unraveling gene-environment interplay to inform Treatment and InterventiON strategies” (ACTION) project. ACTION receives funding from the European Union Seventh Framework Program (FP7/2007-2013) under grant agreement no 602768. Philipp D. Koellinger & Ronald de Vlaming were supported by ERC Consolidator Grant 647648 EdGe. Ian J. Deary, Andrew M. McIntosh, Stuart J. Ritchie, and W. David Hill are members of the University of Edinburgh Centre for Cognitive Ageing and Cognitive Epidemiology, part of the cross council Lifelong Health and Wellbeing Initiative (MR/K026992/1). W. David Hill is supported by a grant from Age UK (Disconnected Mind Project). Polygenic score analyses were conducted under UK Biobank dataset resource-application number 4844.

## Software

GenomicSEM software is an R package that is available from GitHub at the following URL: https://github.com/MichelNivard/GenomicSEM

The GenomicSEM R package can be installed directly at: https://github.com/MichelNivard/GenomicSEM/wiki.

Example GenomicSEM code, including code used to produce results is provided for each set of analyses at the following online wiki: https://github.com/MichelNivard/GenomicSEM/wiki.

## Author Contributions

Software Development: Grotzinger, Rhemtulla, Ip, Nivard, Tucker-Drob

Theory underlying Genomic SEM: Grotzinger, Nivard, Tucker-Drob

Technical Development and Mathematical Derivations: Grotzinger, Rhemtulla, de Vlaming, Nivard, Tucker-Drob

Simulation Studies: Grotzinger, Mallard, Nivard, Tucker-Drob

Polygenic Prediction Analyses: Ritchie, Tucker-Drob

Writing: Grotzinger, Nivard, Tucker-Drob, Feedback and Editing: Rhemtulla, Ritchie, Mallard, Hill, McIntosh, Deary, Keollinger, Harden

## Online Method

### Overview of Genomic SEM

In Genomic SEM, the user specifies a multivariate system of regression and covariance associations involving the genetic components of phenotypes with one another and/or more general latent factors. These associations are represented by parameters that may be fixed or freely estimated, so long as the model is statistically identified (e.g., the number of freely estimated parameters does not exceed the number of nonredundant elements in the genetic covariance matrix being modeled). A set of parameters (*θ*) is estimated such that the fit function indexing the discrepancy between the model-implied covariance matrix ∑ (*θ*) and the empirical covariance matrix *S* is minimized. Model fit is considered good when ∑ (θ) closely approximates *S*.

### Form of Structured Covariance Models

Genomic SEM provides substantial user flexibility with respect to the particular SEM that is specified to produce the model-implied covariance matrix ∑ (*θ*) that approximates the empirical covariance matrix, *S*. SEMs can be partitioned into two sets of equations, one describing the *measurement model*, and the other describing the *structural model*. In the measurement model, the genetic components of *k* “indicator” phenotypes are described as linear functions of a smaller set of *m* (continuous) latent variables, **y = Λη + ε**. In this equation, **y** is a *k* × 1 vector of indicators, **ε** is a *k* × 1 vector of residuals, **η** is an *m* × 1 vector of latent variables, and Λ is a *k* × *m* matrix of factor loadings, i.e. regressions relating the latent variables to the set of indicators. In a typical application of Genomic SEM, each indicator is a function of exactly one of the latent variables (though this so-called “simple structure” restriction may be relaxed). In a confirmatory factor analysis (CFA) model, only the measurement model is specified, and the set of latent variables are allowed to freely covary. Thus, the model-implied covariance matrix of a CFA is **∑** (*θ*) = **ΛΨΛ’**+**Θ**, where **Ψ** is an *m* × *m* latent variable covariance matrix and **Θ** is a *k* × *k* matrix of covariances among the residuals, **ε**. Typically, **Θ** is diagonal, which implies that indicators are mutually independent conditional on the set of latent variables. That constraint may be relaxed such that select pairs of indicators are allowed to covary over and above their associations via the latent variable structure (i.e., residual covariances are allowed). CFA models are typically used to assess the strength of relations between sets of indicators and their respective underlying latent variables, as well as to assess the *fit* of a measurement model to data. A well-fitting CFA model implies that the latent variable structure is able to account for the observed covariances among a set of indicator variables.

When a theory aims to explain associations among latent variables, a structural model can be added to the measurement model to produce a full SEM. The structural model of a SEM relates latent variables to each other via directed regression coefficients. It can be written in matrix notation as **η = Bη + ζ**, where **B** is an *m* × *m* matrix of regression coefficients that relate latent variables to each other and **ζ** is an *m* × 1 vector of latent variable residuals. The model implied covariance matrix of observed variables is **∑** (*θ*) = **Λ (I–B)^-1^ Ψ (I–B’)^-1^ Λ’**+**Θ**, where **I** is an *k* × *k* identity matrix.^30^ Thus, in a full SEM, the empirical matrix is represented by a set of parameters that relate observed variables to latent variables, and relate latent variables to each other in a series of linear equations.

### Path Diagrams

SEMs can be represented graphically as *path diagrams* representing regression and covariance relations among variables.^31^ In path diagrams, observed variables are represented as squares and unobserved (i.e., latent) variables are represented as circles. Regressions relationships between variables are represented as one-headed arrows pointing from the independent variable to the dependent variable. Covariance relationships between variables are represented as two-headed arrows linking the two variables. The variance of a variable (i.e., the covariance between a variable and itself), is represented as a two-headed arrow connecting the variable to itself. In Genomic SEM, we represent the genetic component of each phenotype with a circle, as the genetic component is a latent variable that is not directly measured, but is inferred from LDSC (it is the phenotype itself that is observed in the raw data that is used to produce the summary statistics). SNPs are directly measured, and are therefore represented as squares. When all elements in a SEM are represented and correctly specified in a path diagram, the diagram contains the full system of algebraic equations needed to estimate the full set of SEM parameters, *θ*, and produce the model-implied covariance matrix, ∑ (*θ*).

### Stage 1 Estimation

In Stage 1, the empirical genetic covariance matrix (*S*) and its associated sampling covariance matrix (*Vs*) are estimated using our multivariable extension of LDSC. *S* is a *k* × *k* symmetric matrix with SNP heritabilities on the diagonal and genetic covariances (σ_gi,gj_) between phenotypes *i* and *j* off the diagonal. The genetic covariance between phenotypes *i* and *j* can be computed as the genetic correlation scaled relative to the total genetic variance of each of the two contributing phenotypes (themselves scaled to unit variances), 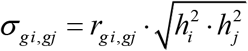. Thus, the genetic covariance matrix of order *k* has *k* = k*(*k +*1)/2 nonredundant elements. It can be written as:

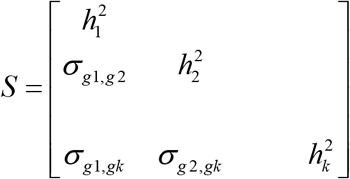

To produce unbiased *SE* estimates and test statistics, we require the asymptotic sampling covariance matrix, *Vs*, of the LDSC estimates that is composed of all nonredundant elements in the *S* matrix. Thus, it is a symmetric matrix of order *k\**. The diagonal elements of *Vs* are sampling variances, that is, squared *SEs* of the elements in *S*. The off-diagonal elements of *Vs* are sampling covariances that indicate the extent to which the sampling distributions of the variance and covariance estimates in *S* covary with one another, as would be expected when there is overlap among the samples from which the terms are estimated.

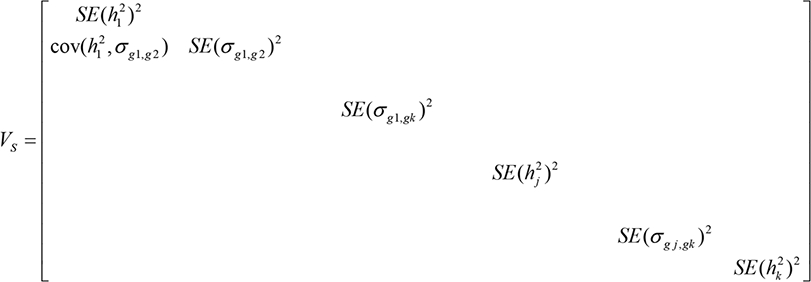

The diagonal elements of *Vs* are estimated using a jackknife resampling procedure in the bivariate version of LDSC that is currently available by its original developers.^4,32^ The LDSC function introduced in the GenomicSEM software package expands the jackknife procedure to the multivariable context in order to produce sampling covariances (which index dependencies among estimation errors) among the elements of *S*, needed to populate the off-diagonal elements of *Vs*.

### Stage 2 Estimation

In Stage 2, the genetic covariance matrix obtained in the previous stage, *S*, is used to estimate the parameters in a SEM. In this stage, we allow for both weighted least squares (WLS) and normal theory maximum likelihood (ML) estimators. WLS does not strictly require positive definite *S* and *Vs* matrices, but may still benefit from positive definiteness during optimization. ML estimation requires both *S* and *Vs* to be positive definite. The GenomicSEM software package therefore smooths *S* and *Vs* to the nearest positive definite matrices prior to Stage 2 estimation using the R function nearPD.^33^

The fit function minimized in the diagonally weighted version of WLS estimation that is standard in the GenomicSEM software package is the following:

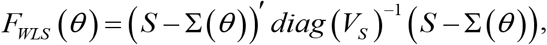

where *diag* (*V_S_*) is the diagonal of the asymptotic covariance matrix, *Vs*. We choose the diagonally weighted version of WLS because it is more tractable to implement for large (highly multivariate) matrices and is more stable than fully weighted WLS in finite samples.^34,35^ We denote the parameter estimates produced in Stage 2 (whether by WLS or ML estimation) as *θ*.

ML estimation proceeds by minimizing the following fit function:

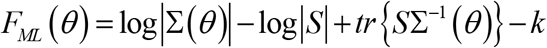

where *∑*(*θ*) is the covariance matrix implied by the set of parameter estimates. Note that, while the formulation of the ML fit function does not explicitly include a weight matrix, it is asymptotically equivalent to a more general formulation that is identical to the WLS fit function, with 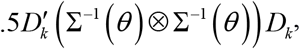, where *D_k_* is the duplication matrix of order *k*, in place of *diag (V_S_)*. Thus, the difference between ML and WLS estimation can be construed as a difference in weight matrices only. A comparison between ML and WLS results can be found in the **Online Supplement** (Supplementary Figs. 18-22, Table S17).

WLS estimation more heavily prioritizes reducing misfit in those cells in the *S* matrix that are estimated with greater precision. This has the desirable property of potentially decreasing sampling variance of the Genomic SEM parameter estimates, which may boost power for SNP discovery and increase polygenic prediction. However, because the precision of cells in the *S* matrix is contingent upon the sample sizes for the contributing univariate GWASs, WLS may produce a solution that is dominated by the patterns of association involving the most well powered GWASs, and contain substantial local misfit in cells of S that are informed by lower powered GWASs. In other words, WLS relative to ML may more heavily prioritize minimizing sampling variance of the parameter estimates in the so-called variance bias tradeoff.36 We expect that this will only occur when the model is overidentified (i.e., *df* > 0), such that exact fit cannot be obtained, and that divergence in WLS and ML estimates will be most pronounced when there is lower sample overlap and the contributing univariate GWASs differ substantially in power. ML estimation may be preferred when the goal is to most evenly weight the contribution of the univariate sample statistics.

Both WLS and ML fit functions will produce consistent estimates of the model parameters when the model is true.^35^ However, the “naïve” *SEs* and fit statistic produced in Stage 2 estimation will be incorrect, because neither estimator uses the full *V_s_* matrix in estimation. Thus, robust corrections must be applied to produce consistent estimates of *SEs* and test statistics. The correct sampling covariance matrix of the Stage Two, Genomic SEM parameter estimates (i.e., *V_θ_*) can be obtained using a sandwich correction:^11,35^

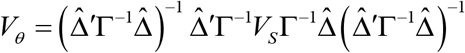

where 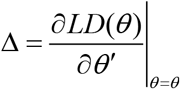 is the matrix of model derivatives evaluated at the two-stage parameter estimates *θ*, Г is the naïve Stage 2 weight matrix that takes its form depending on the estimation method used (WLS or ML), and *Vs* is the sampling covariance matrix of *S* obtained using multivariable LDSC. It may not always be possible to obtain the full sampling covariance matrix, *Vs*. For example, for highly sensitive data only the matrix *S* and the *SEs* of its elements may be available (i.e., the diagonal of *V_s_*). However, we note that when there is low sample overlap across the GWAS’s for each phenotype, off-diagonal elements of the sampling covariance matrix are small and pragmatically ignorable. Moreover, in other contexts with complete sample overlap, *SE* inflation of the SEM parameters estimated using diagonally-weighted versions of WLS has been estimated to be less than 8%^8^ without robustness corrections, and nil with robustness corrections.^35^

### Incorporation of Individual SNP Effects

Several steps are needed to incorporate individual SNP effects into Genomic SEM. The first step requires that the inputted genetic covariance matrix be expanded to include covariances between the SNP and the latent genetic components of each of the phenotypes, g_i_ through g_k_:

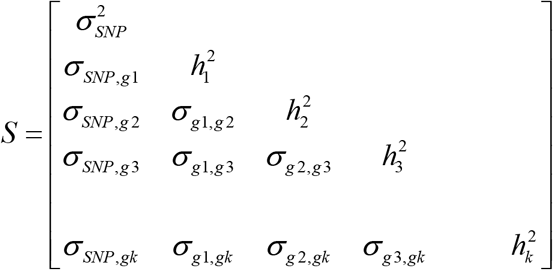

The sampling covariance matrix, *Vs*, associated with this expanded *S* covariance matrix includes a number of components. The sampling variances and sampling covariances of the latent genetic variances (SNP heritabilities) and genetic covariances are obtained from the multivariable LDSC approach introduced above. The SNP variance (derived from reference panel data) is treated as fixed, and its sampling variance and sampling covariance with all other terms are fixed to 0 (or to a very small value to facilitate computational tractability). The sampling covariances of the SNP-genotype covariances with one another are obtained using cross-trait LDSC intercepts (which represent sampling correlations weighted by sample overlap) after being rescaled relative to the sampling variances of the respective SNP-genotype covariances.^9,37^ Finally, the sampling covariance of the SNP-genotype covariances with the genetic variances and genetic covariances are fixed to 0, as sampling variation of the SNP-genotype covariance is expected to be independent of the test statistics of all LD blocks except the one it occupies. Because the sampling variance of the heritabilities and genetic correlations derive from sampling variability in the test statistics within all of the LD blocks, their sampling covariances with a single SNP effect is expected to approach 0.

### Standardization and Scaling of Summary Statistics for Multivariate GWAS

Typically, GWAS summary statistics for quantitative phenotypes are not reported in terms of covariances, but are reported as ordinary least squared (OLS) unstandardized regression coefficients, with the phenotypes standardized prior to analyses (i.e., the coefficients are standardized with respect to the outcome, but not the predictor). In order to transform these partially standardized regression coefficient (b_SNP,P_) of a SNP effect on phenotype P to a covariance, we multiply by the variance of scores on the SNP. The variance 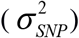 of scores (0, 1, 2) of a biallelic autosomal SNP is estimated as 2×(MAF×(1-MAF)), assuming Hardy-Weinberg-Equilibrium, where MAF is the minor allele frequency, typically obtained from a reference sample. As the latent genetic factors estimated in LDSC are scaled relative to unit-variance scaled phenotypes (by virtue of the SNP heritability estimates being placed on the diagonal of *S*), no further scaling is needed to transform this SNP-phenotype covariance into a SNP-genotype covariance.

When OLS regression coefficients and standard errors are provided from an analysis in which the phenotype has not been standardized prior to analyses, or only Z statistics or *p*-values (for which Z statistics can be readily obtained) are provided, the partially standardized regression coefficients and their standard errors can be obtained as 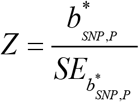, 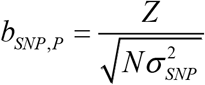 and 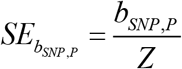, where 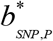 is equal to the regression coefficient for the OLS GWAS of the unstandardized phenotype. These derived partially standardized coefficients are then transformed into covariances by multiplying by the variance of scores on the SNP, as per above.

When the GWAS summary statistics are reported for logistic regressions of liabilities for categorical outcomes (e.g. case/control status) on the SNP, the logistic regression coefficients can be transformed into covariances as above, by multiplying by the SNP variances. However, it is appropriate to further transform the coefficients such that they are scaled relative to unit-variance scaled liability. This can be achieved by dividing by 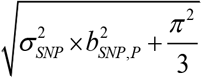, as a logistic regression model implies a residual variance of 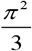. If GWAS summary statistics are reported for odds ratios (ORs), they can be transformed to logistic regression coefficients by taking their natural logarithm. Standard errors for the logistic regression coefficient are obtained as *SE*_OR_/OR. The coefficients and their *SE*s should further be transformed such that they are scaled relative to unit-variance scaled phenotypes, as per above. Note that when the outcomes are categorical, the liability scale heritabilites and genetic covariances from multivariable LDSC (and not what are referred to as the “observed scale” heritabilities and genetic covariances) should be used to populate the *S* matrix. This has the desirable property of both modeling the continuous scale of risk in the population and providing estimates that are independent of the observed prevalence of the categorical outcomes.

On occasion, summary statistics will be provided from OLS GWASs of categorical outcomes (e.g., case/control status). Such an analysis is sometimes referred to as a linear probability model, as it (incorrectly) assumes that the association between the predictor and the probability in being in the comparison (e.g. case) group relative to the reference (e.g. control) group is linear. Parameters from the linear probability model are dependent not only on the strength of the association between the SNP and the continuous underlying liability, but also on the MAF and the proportion of comparison group members (cases) in the sample. Thus, parameters from the linear probability model cannot be used directly in Genomic SEM. However, particularly in the case of complex traits, for which the effect sizes for individual SNPs are small, results from the linear probability model can be used to very closely approximate logistic regression coefficients and standard errors that are amenable for use in Genomic SEM.^38^ The 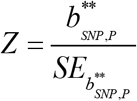, 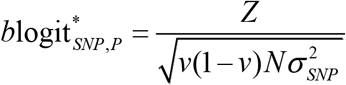, and 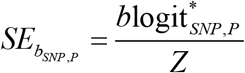, where 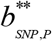 is equal to the regression coefficient from the linear probability model, 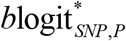 is the expected logistic regression coefficient that is derived from the linear probability model results, *v* is equal to the proportion of cases in the sample, and 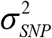 is the variance of the SNP, computed from its MAF obtained from a reference sample, as per above. To scale the derived logistic coefficient such that it is scaled relative to unit-variance scaled liability, the coefficient should be divided by 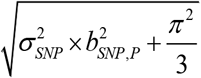. Lloyd-Jones et al. (2018)^38^ report that in a real data analysis of UKB data, the exponentiated regression coefficient (i.e., the odds ratio) obtained directly from a logistic regression-based GWAS and that derived from the linear probability model-based GWAS was nearly perfect (*R*^2^ > 98%, slope ≈ 1). We have verified this nearly perfect correspondence in our own simulations (Supplemental Fig. 23).

### Model Fit Statistics

Model χ^2^ is an index of exact fit of a SEM. It indexes whether the model-implied genetic covariance matrix, ∑(*θ*), differs from the empirical genetic covariance matrix, *S*. Model χ^2^ can also be used as a relative fit index for comparing nested models. Conventional SEM approaches to indexing model χ^2^ are based on formulas that directly incorporate *N*. Because there is not an *N* that directly corresponds to the genetic covariance matrix that is modelled by Genomic SEM in the same way that *N* typically corresponds to an observed covariance matrix, we derived a formula for estimating model χ^2^ that does not require *N*, but instead incorporates the sampling covariance matrix of the model residuals. This is done in two steps. In Step 1, the proposed model (e.g., a common factor model) is estimated. In Step 2, all of the Step 1 estimates are fixed, and the residual covariances and residual variances of the indicators are freely estimated. Residual variances are estimated in Step 2 by estimating the variances of *k* residual factors defined by the indicators. This provides an estimate of the discrepancy between the model implied and observed covariance matrices, 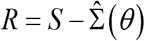, along with the sampling covariance matrix (*V_R_*) of *R*. While the discrepancy between model implied and observed covariance matrices can be computed simply by deriving covariance expectations from the Step 1 model and subtracting the observed covariance matrix, such an approach would not provide the corresponding *V_R_* matrix necessary for the calculations below. The *V_R_* matrix is expected to be positive semidefinite and, consequently, have no negative eigenvalues. Therefore, the *V_R_* matrix has the following eigendecomposition:

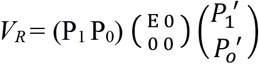

where P_1_ is a matrix of principal components (eigenvectors) of *V_R_*, and *E* is a corresponding diagonal matrix consisting of non-zero eigenvalues. P_0_ reflects the null space of *V_R_*. Projecting *R_i_*—a vector of residual covariances estimated from the Step 2 Model—onto P_1_ and adjusting for corresponding eigenvalues, we have that:

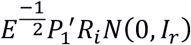

Therefore,

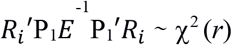

This equation produces a test statistic that is χ^2^ distributed with degrees of freedom (*r*) equal to the difference between the number of nonredundant elements (*k\**) in the empirical covariance matrix (*S*) and the number of freely estimated parameters in the proposed model.

The Comparative Fit Index (CFI) is a test of approximate model fit. CFI indexes the extent to which the proposed model fits better than a model that allows all phenotypes to be heritable, but assumes that they are genetically uncorrelated. The χ^2^ statistic can be used to calculate CFI by calculating a second χ^2^ statistic for a so-called independence model, i.e. a model that estimates genetic variances of all phenotypes but assumes all genetic covariances to be zero, such that ∑(θ) is diagonal. CFI is calculated using the formula below,^39^ with *f* = χ^2^ - degrees of freedom:

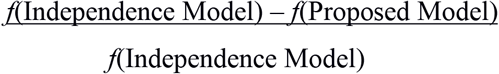

For the χ^2^ of the independence model, a model is estimated in Step 1 that includes only the variance of the indicators and no common factor. In Step 2, these variances are fixed and the covariances among the indicators and variances of *k* residual factors defined by the indicators are estimated and used to populate the same equation above used to calculate the proposed model χ^2^.

Akaike Information Criterion (AIC) is a relative fit index that balances fit with parsimony, and can be used to compare models regardless of whether they are nested. AIC is calculated as:

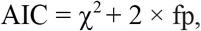

where fp is the number of free parameters in the model.^40^

Standardized Room Mean Square Residual (SRMR) is an index of approximate model fit that is calculated as the standardized root mean squared difference between the model-implied and observed correlations in ∑ (*θ*) and *S*, respectively.^41^ Higher SRMR values indicate a larger discrepancy between ∑(*θ*) and *S*. It is positively-biased, with larger bias resulting when the contributing univariate GWAS samples are lower powered.

#### Validation via Simulation

A generating population with a common factor model defined by four, five, or six indicators was used to examine the null distribution of the newly derived χ^2^ test statistic using a set of 1,0 simulations per model. These simulations did not include individual genotypes, and were simulated solely based on a generating factor structure. For the six indicator models the standardized factor loadings in the generating population were .42, .64, .22, .59, . 19, and .64. The four and five indicator models specified the same factor loadings, excluding the last, or last two loadings, respectively. To allow for potential misfit between the model-implied and observed variances, the variance of the indicators was not standardized. Results indicated that the two-step procedure described above produced a test statistic equivalent to the χ^2^ statistic calculated by lavaan from the raw data (Supplementary Fig. 24 and Table S18). For a χ^2^ distributed test-statistic, the mean of the null sampling distribution should match the *df* of the test. As expected, the distribution of the test-statistic conformed to a χ^2^ distribution with an average approaching the *df* (Supplementary Fig. 25). Calculated CFI values were also highly consistent with those observed using the CFI statistic provided by lavaan when using raw data (Supplementary Fig. 26, Table S18). Calculated AIC values were not contrasted with those obtained using the lavaan package in R in the simulations below as the software uses a formula that includes a log-likelihood estimate contingent on the provided sample size.

### Q_SNP_ Test of Heterogeneity

As with the computation of model χ^2^ outlined above, Q_SNP_ is calculated using a two-step procedure. In Step 1, a *common pathway* model is fit in which both factor loadings, the SNP effect on the common factor(s), and the residual variances of the common and unique factors are freely estimated (with one factor loading fixed to unity for factor identification and scaling). No paths representing direct effects of the SNP on the genetic components of the individual phenotypes are estimated. In Step 2, a *common plus independent pathways model* is specified, in which the factor loadings and the SNP effect on the common factor are fixed to the values estimated in Step 1, and direct effects of the SNP on individual indicators and the residual variances of each indicator are freely estimated. Supplementary Fig. 27 depicts this model, as applied to a single common factor model, with parameters that are fixed in Step 2 depicted in red and those that are freely estimated in Step 2 depicted in black.

#### Null Distribution of Q_SNP_

To verify that the null distribution for Q_SNP_ is χ^2^ distributed, a set of simulations specified a generating population in which the direct effects of the SNP on the indicators were entirely mediated through the common factor (i.e., Q_SNP_ = 0). Each simulation included 1,000 datasets, with *N* = 100,000 completely overlapping participants per dataset. All simulated datasets were analyzed using both WLS and ML. We examined three models with F = 1 factor, and k = 4, 5, or 6 phenotypes. Table S19 presents descriptive statistics for Q_SNP_. Using a genome wide significance threshold, in all cases the false discovery rate for Q_SNP_ was 0, and the power to detect a SNP effect on the common factor was 1. Both WLS and ML estimation produced mean estimates of Q_SNP_ that were approximately equal to the *df* of the corresponding model. Supplementary Fig. 28 depicts the sampling distributions of Q_SNP_ estimated using WLS or ML. Supplementary Fig. 29 plots Q_SNP_ from these two estimation methods against χ^2^ distributions and against one another. These results indicate that both estimation methods produce results that are approximately χ^2^ distributed.

### Genomic SEM Simulation Procedure

In order to evaluate the ability of Genomic SEM to capture the genetic factor structure in the generating population, the GCTA package^3^ was used to generate 100 sets of 6 independent, 100% heritable phenotypes ("orthogonal genotypes") to pair with genotypic data for 39,909 randomly selected, unrelated individuals of European descent from UKB data for the 1,209,498 SNPs present in HapMap3. The generating list of causal SNPs was set to 10,000 for all 600 genotypes, with the specific list of causal variants sampled with replacement from the 1,209,498 SNPs. One of the six orthogonal genotypes per set was designated an index of the general genetic factor and the remaining five were designated indices of domain-specific genetic factors. All of these orthogonal genotypes were scaled to *M*=0, *SD*=1. Five new correlated genotypes were then constructed, each as the weighted linear combination of the general genetic factor and one domain-specific genetic factor. Weights for contribution of the general genetic factor were λ_Fg,k_ =.70, .60, .50, .40, and .30, for correlated genotypes 1-5, respectively. Weights for the domain-specific factors were 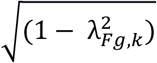. Phenotypes were then each constructed as the weighted linear combination of one of the correlated genotypes and domain-specific environmental factors (randomly sampled from a normal distribution with *M*=0, *SD*=1). Heritabilities for phenotypes 1-5 were set to 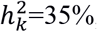, 40%, 50%, 60%, and 70%, respectively, such that the weights for the genotypes were 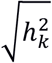 and the weights for the environmental factors were 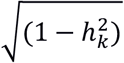. We choose these figures to stabilize the properties of the distributions across simulations at 100 replications with *N*~39K each. We expect that with lower SNP h^2^’s, the same patterns would hold, albeit at larger sample sizes. Each of the 500 phenotypes (100 sets of 5 phenotypes) was then analyzed as a univariate GWAS in PLINK^42^ to produce univariate GWAS summary statistics. Our multivariable LDSC function was then used to construct 100 sets of 5×5 genetic covariance matrices (*S*) and associated sampling covariance matrices (*Vs*), and Genomic SEM was used to fit a one factor model to each set.

### Quality Control Procedures

#### LD-Score Regression

For the *p*-factor, neuroticism, and anthropometric traits, quality control (QC) procedures for producing the *S* and *Vs* matrix followed the defaults in LDSC. We recommend using these defaults for multivariable LDSC, including removing SNPs with an MAF < 1%, information scores < .9, SNPs from the MHC region, and filtering SNPs to HapMap3. Quality control procedures for the multivariable regression example mirrored those used by Nieuwboer et al. (2016)^21^ for comparative purposes. More specifically, SNPs were excluded with MAFs < .05 as determined by the HapMap Consortium,^43^ and with information values less than 0.9 or greater than 1.1. SNPs were also filtered to HapMap3.

#### Multivariate GWAS

To obtain summary statistics for multivariate GWAS, we recommend using QC procedures of removing SNPs with an MAF < .01 in the reference panel, and those SNPs with an INFO score < 0.6. MAFs were obtained for the current analyses using the 1000 Genomes Phase 3 reference panel. Using these QC steps, 1,979,881 SNPs were present across schizophrenia, bipolar disorder, MDD, PTSD, and anxiety. For neuroticism, there were 7,265,104 SNPs that were present across all phenotypes. These QC procedures are the defaults for the processing function within the GenomicSEM package. The regression effects for the univariate indicators of the *p*-factor were standardized using the procedure for logistic coefficients outlined above. Regression effects for neuroticism indicators were converted from linear probability to logistic coefficients and then standardized with respect to the variance in the outcome.

### Out-of-Sample Prediction

Genomic SEM analyses that were used to produce the summary statistics for construction of polygenic scores for out-of-sample prediction omit the PGHC MDD 2018 GWAS and SCZ 2018 GWAS and replace them with the PGC MDD 2013^44^ and PGC SCZ 2014^45^ GWAS to prevent overlap between discovery and target samples. This resulted in a Genomic SEM-based multivariate GWAS using 930,581 SNPs. Analyses used to construct a phenotypic *p*-factor for polygenic prediction in the UKB dataset were restricted to data on up to *N*=332,050 European participants. The Genomic SEM of the *p*-factor employed case-control GWAS statistics to construct summary statistics for a general factor of liability for clinically-severe levels of psychopathology as the discovery phenotype. For out-of-sample prediction, we selected a set of psychiatric symptoms (rather than diagnoses) to construct liability for general and domain-specific factors of psychiatric symptomology across the subclinical-to-clinical ranges as the target phenotypes. From the UKB dataset, we chose symptoms falling within the following domains: psychosis, mania, depression, post-traumatic stress, and anxiety. We fit a confirmatory factor model (diagram shown in Supplementary Fig. 29) to the phenotypic symptom endorsements, treating them as ordered categorical variables. Analyses were run in M*plus*,^46^ with the target phenotypes—the *p*-factor and each of the individual domains—specified as latent variables. To construct PGSs, we removed from both the *p*-factor and univariate summary statistics the 5 SNPs that were identified as having genome-wide significant Q_SNP_ estimates for ML, along with SNPs that were in LD with these SNPs using an *r*^2^ threshold of 0.1 and 500kb window. PGSs were constructed using PRSice,^47^ with LD clumping was set to *r*^2^ > 0.25 over 250kb sliding windows. PGSs for the *p*-factor were based on the WLS and ML summary statistics produced using Genomic SEM. We ran PGS analyses using a *p*-value threshold of 1.0 (i.e., we used all available SNPs apart from those removed due to Q_SNP_ analyses). In order to maintain comparability, PGSs for the univariate summary statistics were constructed based on the same SNPs with which the PGSs for the *p*-factor were constructed. In the confirmatory factor models, we included controls for age, sex, genotyping array, and 40 principal components of ancestry in conjunction with the PGS predictor.

### Clumping and Biological Annotation

Lead SNPs for univariate indicators and the common factors were identified using the clumping algorithm in PLINK^42^ We defined LD-independent SNPs using an *r*^2^ threshold of 0.1 and a 500-kb window using the same 1000 Genomes Phase 3 reference panel used for obtaining MAF. For chromosomes 6 and 8 an additional pruning filter was used of 1,000,000 kb and *r*^2^ > 0.1 to account for long-range LD due to the MHC region and pericentric inversion, respectively. The lead SNPs identified using PLINK were entered into DEPICT. Prioritized genes, enriched gene sets, and enriched tissues were identified using the standard false discovery rate of 5%.

### Description of GenomicSEM Software

The Genomic SEM software package, GenomicSEM, is written as an R package and is available through GitHub at https://github.com/MichelNivard/GenomicSEM. GenomicSEM contains several functions, including procedures for QCing and standardizing summary statistics, a function for producing genetic covariance matrices (*S_LD_*) and their associated sampling covariance matrices (*V_LD_*) using a multivariable extension of LD Score regression, functions for fitting Genomic Structural Equation Models to *S* and *Vs*, and functions for adding SNP level data to the *S* and *Vs* matrices that are used for implementing Genomic SEM for multivariate GWAS discovery. Functions include both pre-specified models (e.g., a single common factor model) and user-specified models. Output includes both unstandardized and standardized solutions, along with the fit indices described above. WLS estimation is the default in the GenomicSEM package. GenomicSEM uses the lavaan Structural Equation Modeling package^48^ as the primary workhorse for model specification and numerical optimization. We also provide limited support for OpenMx^49^ To run the multivariable LDSC function on five phenotypes takes ~15 minutes. For models of multivariate genetic architecture that do not incorporate individual SNP effects, the typical run time is ~1 second on a laptop. Using parallel processing implemented in the GenomicSEM package on an 8-core laptop, a multivariate Genomic SEM GWAS with five indicators and ~1 million SNPs took ~8 hours.

## References

1. Lee S. H. et al. Genetic relationship between five psychiatric disorders estimated from genome-wide SNPs. Nature Genetics 45, 984 (2013).

2. Bush, W. S., Oetjens, M. T. & Crawford, D. C. Unravelling the human genome-phenome relationship using phenome-wide association studies. Nature Reviews Genetics 17, 129 (2016).

3. Yang, J., Lee, S. H., Goddard, M. E. & Visscher, P. M. GCTA: a tool for genome-wide complex trait analysis. The American Journal of Human Genetics 88, 76–82 (2011).

4. ReproGen Consortium et al. An atlas of genetic correlations across human diseases and traits. Nature Genetics 47, 1236–1241 (2015).

5. Day F. R. et al. Shared genetic aetiology of puberty timing between sexes and with health-related outcomes. Nat Comms 6, 8842 (2015).

6. McLaughlin R. L. et al. Genetic correlation between amyotrophic lateral sclerosis and schizophrenia. Nat Comms 8, 14774 (2017).

7. Zipfel, S., Giel, K. E., Bulik, C. M., Hay, P. & Schmidt, U. Anorexia nervosa: aetiology, assessment, and treatment. The Lancet Psychiatry 2, 1099–1111 (2015).

8. Verhulst, B., Maes, H. H. & Neale, M. C. GW-SEM: A Statistical Package to Conduct Genome-Wide Structural Equation Modeling. Behav Genet 47, 345–359 (2017).

9. Turley P. et al. Multi-trait analysis of genome-wide association summary statistics using MTAG. Nature Genetics 1 (2018).

10. Cheung M. W.-L. metaSEM: An R package for meta-analysis using structural equation modeling. Frontiers in Psychology 5, 1521 (2015).

11. Savalei V. & Bentler P. M. A two-stage approach to missing data: Theory and application to auxiliary variables. Structural Equation Modeling: A Multidisciplinary Journal 16, 477–497 (2009).

12. Yuan K. H. & Bentler P. M. Robust mean and covariance structure analysis through iteratively reweighted least squares. Psychometrika 65, 43–58 (2000).

13. de Vlaming, R., Johannesson, M., Magnusson, P. K., Ikram, M. A. & Visscher, P. M. Equivalence of LD-score regression and individual-level-data methods. bioRxiv 211821 (2017).

14. Lee J. J. & Chow C. C. LD Score regression as an estimator of confounding and genetic correlations in genome-wide association studies. (2017). doi: 10.1101/234815

15. Browne M. W. Asymptotically distribution-free methods for the analysis of covariance structures. British Journal of Mathematical and Statistical Psychology 37, 62–83 (1984).

16. Huedo-Medina, T. B., Sánchez-Meca, J., Marín-Martínez, F. & Botella, J. Assessing heterogeneity in meta-analysis: Q statistic or I^2^ index? Psychological Methods 11, 193 (2006).

17. Caspi A. et al. The p Factor. Clinical Psychological Science 2, 119–137 (2013).

18. Stochl J. et al. Mood, anxiety and psychotic phenomena measure a common psychopathological factor. Psychological Medicine 45, 1483–1493 (2015).

19. Pettersson, E., Larsson, H. & Lichtenstein, P. Common psychiatric disorders share the same genetic origin: a multivariate sibling study of the Swedish population. Molecular Psychiatry 21, 717–721 (2015).

20. Seed C. et al. Hail: An Open-Source Framework for Scalable Genetic Data.

21. Nieuwboer, H. A., Pool, R., Dolan, C. V., Boomsma, D. I. & Nivard, M. G. GWIS: Genome-Wide Inferred Statistics for Functions of Multiple Phenotypes. The American Journal of Human Genetics 99, 917–927 (2016).

22. Rietveld C. A. et al. GWAS of 126,559 individuals identifies genetic variants associated with educational attainment. science 1235488 (2013).

23. Ruderfer D. M. et al. Cross-Disorder Working Group of Psychiatric Genomics Consortium (2014). Polygenic dissection of diagnosis and clinical dimensions of bipolar disorder and schizophrenia. Mol. Psychiatry 19, 1017–1024

24. Rhemtulla, M., Brosseau-Liard, P. É. & Savalei, V. When can categorical variables be treated as continuous? A comparison of robust continuous and categorical SEM estimation methods under suboptimal conditions. Psychological Methods 17, 354–373 (2012).

25. Pers T. H. et al. Biological interpretation of genome-wide association studies using predicted gene functions. Nat Comms 6, 5890 (2015).

26. Li Z. et al. Genome-wide association analysis identifies 30 new susceptibility loci for schizophrenia. Nature Genetics 49, 1576 (2017).

27. Hu Y. et al. GWAS of 89,283 individuals identifies genetic variants associated with self-reporting of being a morning person. Nat Comms 7, 10448 (2016).

28. Consortium A. S. D. W. G. O. T. P. G. et al. Meta-analysis of GWAS of over 16,000 individuals with autism spectrum disorder highlights a novel locus at 10q24. 32 and a significant overlap with schizophrenia. Molecular autism 8, 1–17 (2017).

29. Hill W. D. et al. A combined analysis of genetically correlated traits identifies 187 loci and a role for neurogenesis and myelination in intelligence. Molecular Psychiatry 1 (2018).

## Online Methods-only References

30. Jöreskog K. G. & Sörbom D. LISREL 8: Structural equation modeling with the SIMPLIS command language. (1993).

31. Boker S. M. & McArdle J. J. Path analysis and path diagrams. Wiley StatsRef: Statistics Reference Online (2014).

32. Bulik-Sullivan B. K. et al. LD Score regression distinguishes confounding from polygenicity in genome-wide association studies. Nature Genetics 47, 291 (2015).

33. Sparse and Dense Matrix Classes and Methods [R package Matrix version 1.2-12]. (2017).

34. Flora D. B. & Curran P. J. An empirical evaluation of alternative methods of estimation for confirmatory factor analysis with ordinal data. Psychological Methods 9, 466 (2004).

35. Savalei V. Understanding robust corrections in structural equation modeling. Structural Equation Modeling: A Multidisciplinary Journal 21, 149–160 (2014).

36. Yarkoni T. & Westfall J. Choosing prediction over explanation in psychology: Lessons from machine learning. Perspectives on Psychological Science 12, 1100–1122 (2017).

37. Baselmans B. M. et al. Multivariate Genome-wide and integrated transcriptome and epigenome-wide analyses of the Well-being spectrum. bioRxiv 115915 (2017).

38. Lloyd-Jones, L. R., Robinson, M. R., Yang, J. & Visscher, P. M. Transformation of summary statistics from linear mixed model association on all-or-none traits to odds ratio. Genetics genetics. 300360.2017 (2018).

39. Kenny D. A. Measuring model fit. (2015). http://davidakenny.net/cm/fit.htm

40. Tanaka J. S. Multifaceted conceptions of fit in structural equation models. Sage focus editions 154, 10–10 (1993).

41. Hu L. T. & Bentler P. M. Cutoff criteria for fit indexes in covariance structure analysis: Conventional criteria versus new alternatives. Structural Equation Modeling: A Multidisciplinary Journal 6, 1–55 (1999).

42. Purcell S. et al. PLINK: a tool set for whole-genome association and population-based linkage analyses. The American Journal of Human Genetics 81, 559–575 (2007).

43. Consortium I. H. The international HapMap project. Nature 426, 789 (2003).

44. Ripke S. et al. A mega-analysis of genome-wide association studies for major depressive disorder. Molecular Psychiatry 18, 497 (2013).

45. Ripke S. et al. Biological insights from 108 schizophrenia-associated genetic loci. Nature 511, 421 (2014).

46. Muthen L. K. & Muthen B. O. Mplus: The comprehensive modeling program for applied researchers [Computer program]. (Los Angeles: Muthen & Muthen, 1998).

47. Euesden, J., Lewis, C. M. & O'Reilly, P. F. PRSice: polygenic risk score software. Bioinformatics 31, 1466–1468 (2014).

48. Rossel Y. Lavaan: An R package for structural equation modeling and more. Version 0.5-12 (BETA). Retrieved from http://users._ugent._be/~yrosseel/lavaan/lavaanlntroduction.pdf (2012).

49. Neale M. C. et al. OpenMx 2.0: Extended structural equation and statistical modeling. Psychometrika 81, 535–549 (2016).

